# Engineering the AEG1 promoter from cotton to develop male sterile lines

**DOI:** 10.1101/2021.03.18.435223

**Authors:** Kamlesh Kumar Soni, Amita Kush Mehrotra, Pradeep Kumar Burma

## Abstract

This work reports on modifying the Upstream Regulatory Module (URM, 1.5 Kb region upstream of the open reading frame) of *Anther Expressing Gene 1* (*AEG1*) from cotton to achieve anther specific activity. *AEG1* was identified in a previous study aimed to isolate a promoter with tapetum specific activity. Such a promoter could then be used to express *barnase* and *barstar* genes for developing male sterile and restorer lines for hybrid seed production in cotton. The AEG1 URM was observed to be active in tapetum as well as in roots making it unusable to drive the expression of *barnase* gene. Analysis of the URM showed the presence of several root specific motifs. Two modified AEG1 URMs were developed, by removing or mutating these motifs and its activity checked in tobacco. The activity of one of the modified URMs, AEG1(ΔBmut) was restricted to the anther tissue as observed using the reporter gene *ß-glucuronidase*. The study also demonstrates that male sterile lines could be developed in tobacco using the AEG1(ΔBmut) URM to express the *barnase* gene. This work thus shows the possibility of engineering promoters to achieve tissue specificity and to develop male sterile lines in cotton.

---

Tapetum specific promoters are key to develop *barnase* and *barstar* gene based male sterile and restorer lines for production of hybrid seeds (Mariani *et al*., 1992). The *Nicotiana tabacum* (tobacco) TA29 promoter has been used to develop male sterile and restorer lines in crops like Brassica (Jagannath *et al*., 2001, 2002). In order to expand the technology for cotton, our laboratory identified the Anther Expressing Gene 1 (*AEG1*,Paritosh *et al*., 2018), which from a microarray-based comparative analysis of the transcriptome from different tissues was found to express in anthers from *Gossypium hirsutum. In situ* hybridization showed presence of *AEG1* transcripts in the tapetum. Cotton transgenic lines carrying 1.5 Kb region upstream of the open reading frame (URM, Upstream Regulatory Module) to drive an intron containing *ß-glucuronidase* gene (Gi) as reporter confirmed GUS activity in tapetum. However, GUS activity was also observed in roots. This made the URM unusable to drive *barnase* gene, which expresses a cytotoxic protein and, therefore, expression in roots would be highly undesirable. Analysis of the 1.5 Kb URM for presence of different *cis*-elements showed that apart from elements like POLLENLAT52 and GTGANT10 observed in genes expressed in anthers, it also contained multiple copies of the *cis*-element, ROOTMOTIFTAPOX1. This root motif with a consensus sequence of ‘ATATT’ is observed in the rolD promoter of *Agrobacterium rhizogenes*, which is strongly expressed in tobacco roots (Elmayan & Tepfer, 1995). It is also present in promoter of the root specific wheat peroxidase (Elmayan & Tepfer, 1995) and Arabidopsis glycosyltransferase genes (Vijaybhaskar *et al*., 2008). A total of 12 motifs were observed, nine of which were clustered between −285 and −452 bp region and three at −53, −97 and −237 positions of the URM (Figure 1a).

**Figure 1.**
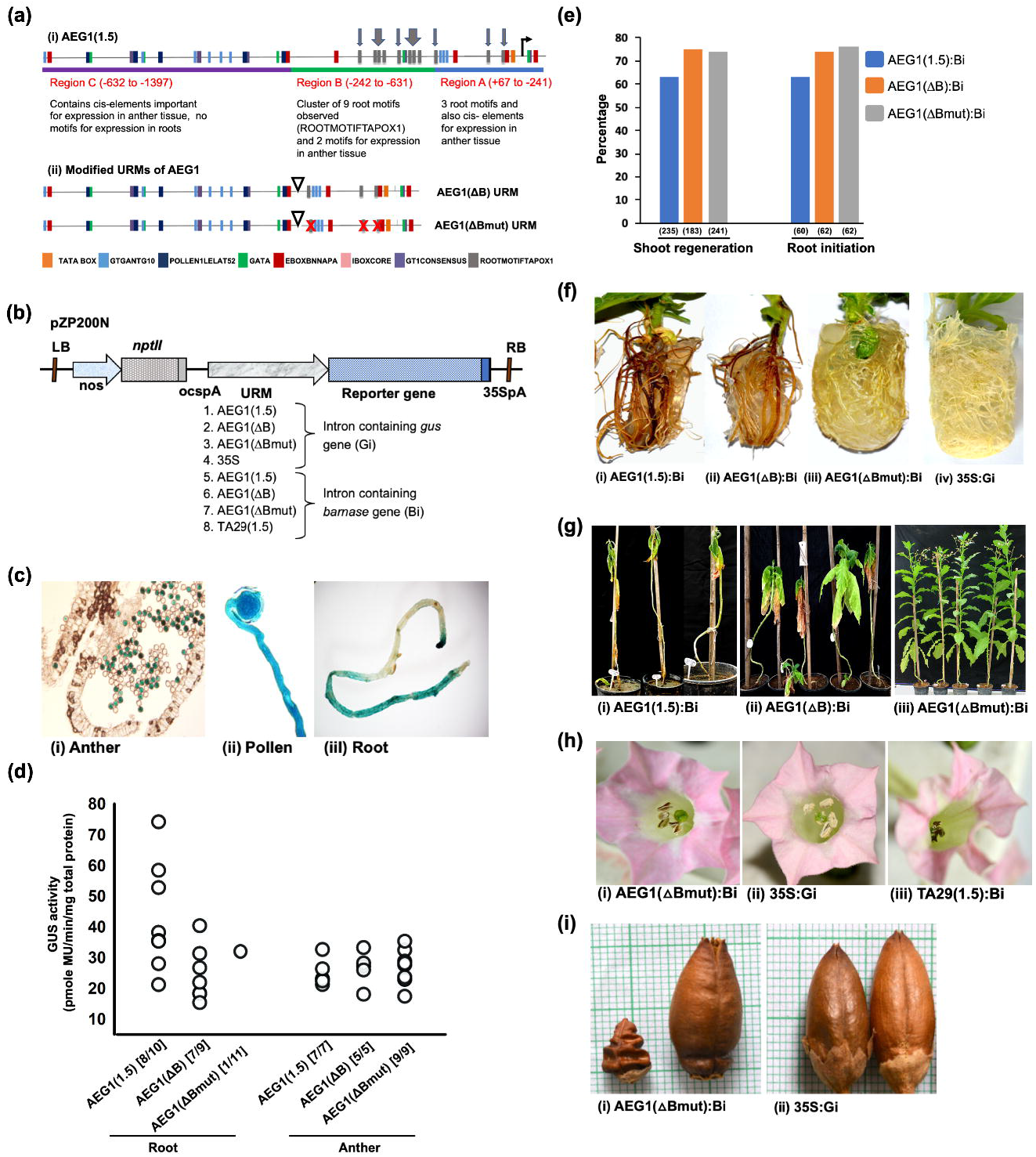
(a) Organization of AEG1 URMs. Cis-elements observed in wild type URM 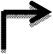 and modified URMs. [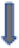 root motifs, transcription start site, 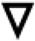 deletion and 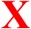 mutation]. (b) Outline of binary vectors used for transformation of tobacco. (c) GUS staining in anthers, germinating pollen and roots of AEG1(1.5) URM transgenic lines. (d) GUS activity driven by different URMs in transgenic lines (number of lines showing recordable activity out of the total in parenthesis). (e) Percent of explants transformed with AEG1URM:Bi constructs forming shoots and shoots developing roots (number of explants or shoots examined is in parentheses). (f, g) Phenotype of roots in tissue culture and plants in greenhouse, respectively of transgenic lines developed with different AEG1URM:Bi and 35S:Gi constructs. (h) Male sterile flowers in lines developed with AEG1(ΔBmut):Bi and TA29(1.5):Bi constructs compared to male fertile lines with 35S:Gi. (i) Phenotype of seed capsules in lines with AEG1(ΔBmut):Bi and 35S:Gi. The capsule on left is following self-pollination while that on the right is after crosspollination.

Here, we engineered the wild type AEG1 URM [called AEG1(1.5)] to make its expression tapetum specific. We developed two different URMs: (i) AEG1(ΔB), wherein the cluster of 9 root motifs in the −285 and −452 bp region were deleted by removing a DNA fragment flanked by *Pac*I sites at −242 and −631 bp and (ii) AEG1(ΔBmut), wherein the 3 proximal root motifs were mutated in the background of AEG1(ΔB), thus removing all the root motifs in AEG1(1.5) URM (Figure 1a). The activity of these URMs was studied in transgenic tobacco lines using the Gi reporter or an intron containing *barnase* gene (Bi). Tobacco, rather than cotton was used because of ease in generating multiple transgenic lines in a shorter time. Further, Bi was used a reporter in addition to Gi as (i) the *barnase* gene gives a more sensitive read-out for tissue specificity (Sharma *et al*.,2018) and (ii) would allow testing the possibility of developing male sterile lines using the re-engineered URMs. Constructs having Gi under the control of CaMV 35S promoter (35S:Gi) and Bi under the control of tapetum specific promoter TA29 (TA29:Bi). were used as controls. Eight different expression cassettes were developed in the binary vector pPZP200N (Figure 1b). Following *Agrobacterium*-mediated transformation the activity of the URM in the URM:Gi lines was monitored through histochemical and biochemical assay of the GUS activity. Histochemical staining of leaf and stem of independent transgenic lines showed no activity of the URM in these tissues while ~76% of the 17 independent AEG1(1.5):Gi cassette carrying lines (N = 17) showed GUS expression in roots. In lines carrying AEG1(ΔB):Gi, ~66% of the lines showed expression in roots. On the other hand, expression in roots was substantially reduced to ~19% in case of lines (N = 26) carrying AEG1(ΔBmut):Gi. Staining of anthers for GUS activity showed that wild type URM [AEG1(1.5)] expressed in the pollen (Figure 1c). and not in the tapetum tissue as observed in case of cotton. Quantification of GUS activity in root showed that there was an overall reduction in the activity of the AEG1(ΔB) (Figure 1d), while no recordable GUS activity was observed in 10 out of the 11 lines analyzed with AEG1(ΔBmut). In case of anthers, spread of GUS activity was similar between the three URM:Gi lines. These observations showed that the URM, AEG1(ΔBmut) retains its expression in the anther while not expressing in roots in majority of the transgenic lines.

Transformation of tobacco was also carried out with the above noted URM:Bi constructs as well as 35S:Gi and TA29:Bi as controls. Regeneration was monitored for shoot formation from explants and rooting to assess any activity of the URMs during the process of transformation and regeneration. Expression of the *barnase* gene in any vegetative tissue would adversely affect the regeneration process. As shown in Figure 1e there was no major difference in the frequency of shoot and root regeneration in the lines developed with the different URM:Bi constructs. However, the roots of lines developed with the constructs AEG1(1.5):Bi and AEG1(ΔB):Bi were adversely affected. The roots were of low density, which became brown and shriveled in few days. On the other hand, roots of 16 out of 20 of AEG1(ΔBmut):Bi lines showed healthy architecture comparable to the 35S:Gi lines (Figure 1f). On transfer to soil, only the AEG1(ΔBmut):Bi lines survived with proper vegetative growth comparable to the control lines (Figure 1g). On flowering, 9 of the 11 lines transferred were male sterile (Figure 1h), also reflected in lack of seed formation and shriveled capsule on self-pollination (Figure 1i). The remaining 2 lines had pollens and set seed on pollination. The occurrence of a few male fertile lines among lines expected to be sterile is not uncommon and has been reported in lines developed with TA29 promoter controlled *barnase* gene (Jagannath *et al*., 2001). The observed male sterility was surprising since the GUS staining pattern would suggest only reduced pollens as the *barnase* gene would have segregated following meiosis. Therefore, male sterility indicates that the AEG1 URM is active at earlier microsporogenesis stages, which was not detected by GUS activity. All male sterile lines set seeds on cross pollination with untransformed tobacco lines showing that there was no leaky activity of the AEG1(ΔBmut) in the female parts (Figure 1i). The seeds from cross-pollinated lines germinated at frequency ranging from 76 to 81%, which was similar to seeds from plants developed with 35S:Gi construct. Segregation of the kanamycin resistant and sensitive phenotypes in progeny of cross-pollinated lines showed that most of the lines had integration at a single locus, another hallmark of male transgenic lines developed with *barnase* gene.

To the best of our knowledge there are no reports in plants where such engineering of promoter has been done to achieve tissue specificity, although there are several reports on developing synthetic promoters (reviewed by Ali & Kim, 2019). Literature survey showed one report involving human cell lines where in a constitutive gene was modified to make it cell type specific (Kuzmin *et al*., 2010).

There are several reports of promoters or genes being active/ expressed in anther tissue of cotton e.g. *GhACS1* (Wang and Li, 2009) but none of them have been critically checked for its specificity which is important for use in the *barnase - barstar* system. The engineered AEG1(ΔBmut) URM thus opens up the possibility to develop male sterile lines in cotton using the *barnase* gene.

## Acknowledgements

The work was supported by R&D projects of University of Delhi and grant-in-aids to the department from UGC and DST, Govt. of India. KKS and AKM were supported by fellowships from UGC and CSIR, respectively. The authors thank Prof. S C Lakhotia for his inputs.

## Contributions

PKB conceptualized the project. PKB, KKS and AKM designed experiments. AKM and KKS carried out experiments. PKB, AKM and KKS analyzed the data. PKB and KKS wrote the manuscript.

## References

1. Ali, S., & Kim, W. C. (2019) A Fruitful Decade Using Synthetic Promoters in the Improvement of Transgenic Plants. Front. Plant Sci. 10, 1433.

2. Elmayan, T., Tepfer, M. (1995) Evaluation in tobacco of the organ specificity and strength of the *rolD* promoter, domain A of the 35S promoter and the 35S^2^ promoter. Transgenic Res. 4, 388–396.

3. Jagannath, A., Arumugam, N., Gupta, V., Pradhan, A., Burma, P., & Pental, D. (2002) Development of transgenic barstar lines and identification of a male sterile (barnase)/ restorer (barstar) combination for heterosis breeding in Indian oilseed mustard (Brassica juncea). Current Science 82 (1), 46–52.

4. Jagannath, A., Bandyopadhyay, P., Arumugam, N., Gupta, V., Burma, P. K., Pental, D. (2001) The use of a Spacer DNA fragment insulates the tissue-specific expression of a cytotoxic gene (*barnase*) and allow high-frequency generation of transgenic male sterile lines in Brassica juncea L. Mol. Breeding 8, 11–23.

5. Kuzmin, D., Gogvadze, E., Kholodenko, R., Grezla D.P., Mityaev, M., Vinogradova, T. et al. (2010) Novel strong tissue specific promoter for gene expression in human germ cells. BMC Biotechnol. 10, 58.

6. Mariani, C., Gossele, V., Beuckeleer, M.D., Block, M. D., Goldberg, R.B., Greef, W.D., Leemans J (1992) A chimeric ribonuclease-inhibitor gene restores fertility to male sterile plants. Nature 357, 384–387.

7. Paritosh, K., Singh, A.K., Mehrotra, A.K., Pental D., Burma, P.K. (2018) Identification and characterization of the promoter of a gene expressing mainly in the tapetum tissue of cotton (Gossypium hirsutum L.). Plant Biotechnol. Rep. 12, 377–388.

8. Sharma, P.A., Verma, N., Burma, P.K. (2018) Analysis of activity driven by upstream regulatory modules (URM) of tapetum specific genes TA29 and A9 at ectopic locations in tobacco transgenics. J. Plant Biochem Biotechnol. 27, 443–452

9. Vijaybhaskar, V., Subbiah, V., Kaur, J., Vijayakumari, P., Siddiqi, I. (2008) Identification of a root-specific glycosyltransferase from *Arabidopsis* and charcaterization of its promoter. J. Biosci. 33(2), 185–193

10. Wang, X. L., & Li, X. B. (2009) The GhACS1 gene encodes an acyl-CoA synthetase which is essential for normal microsporogenesis in early anther development of cotton. Plant J. 57(3), 473–486.

